# Loss-of-function of *Fbxo10*, encoding a post-translational regulator of BCL2 in lymphomas, has no discernible effect on BCL2 or B lymphocyte accumulation in mice

**DOI:** 10.1101/2020.08.05.237545

**Authors:** Etienne Masle-Farquhar, Amanda Russell, Yangguang Li, Fen Zhu, Lixin Rui, Robert Brink, Christopher C Goodnow

**Affiliations:** Immunology Division, Garvan Institute for Medical Research, Sydney, NSW, Australia; Division of Hematology/Oncology, Department of Medicine, University of Wisconsin, 4060 WIMR, 1111 Highland Ave, Madison, WI 53705; St Vincent’s Clinical School, University of New South Wales, Sydney, NSW, Australia; School of Medical Sciences and Cellular Genomics Futures Institute, UNSW Sydney, Sydney, NSW, Australia

## Abstract

Regulation of the anti-apoptotic BCL2 protein determines cell survival and is frequently abnormal in B cell lymphomas. An evolutionarily conserved post-translational mechanism for over-expression of BCL2 in human B cell lymphomas and the BCL2 paralogue CED-9 in *Caenorhabditis elegans* results from loss-of-function mutations in human FBXO10 and its *C.elegans* paralogue DRE-1, a BCL2/CED-9-binding subunit of the SKP-CULLIN-FBOX (SCF) ubiquitin ligase. Here, we tested the role of FBXO10 in BCL2 regulation by producing mice with two different CRISPR/*Cas9*-engineered *Fbxo10* mutations: an Asp54Lys (E54K) missense mutation in the FBOX domain and a Cys55SerfsTer55 frameshift (fs) truncating mutation. Mice homozygous for either mutant allele were born at the expected Mendelian frequency and appeared normal in body weight and appearance as adults. Spleen B cells from homozygous mutant mice did not have increased BCL2 protein, nor were the numbers of mature B cells or germinal centre B cells increased as would be expected if BCL2 was increased. Other lymphocyte subsets that are also regulated by BCL2 levels also displayed no difference in frequency in homozygous *Fbxo10* mutant mice. These results support one of two conclusions: either FBXO10 does not regulate BCL2 in mice, or it does so redundantly with other ubiquitin ligase complexes. Possible candidates for the latter include FBXO11 or ARTS-XIAP. The difference between the role of FBXO10 in regulating BCL2 protein levels in *C. elegans* and in human DLBCL, relative to single-gene deficient mouse leukocytes, should be further investigated.

## Introduction

Survival of many cells, notably mature B-lymphocytes, is promoted by and depends upon the *Bcl2* gene encoding an essential inhibitor of apoptosis [1–5]. The *B-cell leukemia-lymphoma-2* (*BCL2*) gene was discovered because hybrid *BCL2-Immunoglobulin Heavy chain* (*IGH*) fusion transcripts [6–8] resulting in aberrantly high BCL2 protein expression [9] are often created by a t(14; 18) chromosomal translocation that occurs in 85% of human follicular B cell lymphomas [10, 11] and 34% of germinal centre (GC)-type diffuse large B-cell lymphomas (DLBCL) [12]. While expressed in other mature B cell subsets, BCL2 is absent in normal GC B cells due to BCL6-mediated transcriptional suppression [13, 14], but this regulation is disrupted by t(14;18) that brings *BCL2* under control of the constitutively active *IGH* promoter [12, 15]. *BCL2* over-expression due to 18q21 amplification or activated NF-κB signalling often occurs in activated B cell (ABC)-type DLBCL [16]. Missense *BCL2* point mutations are also frequently observed, associated with activation-induced cytidine deaminase (AID)-mediated somatic hypermutation (SHM) and exhibiting significant negative selection against *BCL2* loss-of-function mutations [17]. Together, translocations, amplifications and missense mutations make *BCL2* the second most highly mutated gene in DLBCL [18].

BCL2 is a moderately long-lived protein with a 10-hour half-life in mature B cells [19, 20]. Stability of BCL2 and its anti-apoptotic paralogues, relative to the even longer-lived pro-apoptotic BAX and BAK proteins, is a key determinant of anti-apoptotic potency [21, 22]. Despite the importance of BCL2 regulation for normal and neoplastic lymphocytes, remarkably little is known about mechanisms controlling BCL2 protein accumulation and turnover [23]. Protein ubiquitination resulting in proteasomal degradation is an important mechanism determining protein stability. An important family of protein ubiquitin ligases comprise the S phase kinase-associated protein 1 (SKP1)–cullin 1 (CUL1)–F-box protein (SCF) complexes [24, 25]. Specific protein substrates for ubiquitination by a given SCF complex are recognised by diverse domains in the 69 different FBOX proteins. The FBOX domain itself mediates interaction with SKP1, which in turn binds CUL1 and RBX1 to activate the E2 ubiquitin ligase.

In a genetic screen in *Caenorhabditis elegans*, Chiorazzi *et al.* [26] identified a strain with a recessive S275L missense mutation in the F-box domain of the product of gene *dre-1* that prevented apoptosis of the tail spike cell. Two additional *dre-1* alleles had a similar effect and complementation confirmed the variant as causal, whilst transgenic over-expression of *dre-1* resulted in an opposing effect of increased apoptosis. The DRE-1 protein bound weakly to the *C. elegans* BCL2 homologue, CED-9, and strong epistasis occurred between a weak loss-of-function *dre-1* mutation and a weak loss-of-function *Ced-9* mutation. The phenotypic effects of the *dre-1* mutation were recapitulated by RNA interference (RNAi) against C. *elegans* SKP-Cullin complex proteins *skr-1* and *cul-1*, and expression followed by co-immunoprecipitation showed the *dre-1* S275L FBOX domain mutation diminished DRE-1 binding to the C.elegans SKP1 paralogue.

The C.elegans DRE-1 protein most closely resembles two human proteins, FBXO11 and FBXO10, with FBXO11 being the closest homologue [27]. Only FBXO10 and FBXO11 have the same combination of F-box and a Carbohydrate-binding proteins And Sugar Hydrolases (CASH) domain as DRE-1, but FBOX11 is confined to the nucleus where it controls BCL6 protein levels [28] whereas BCL2 is cytoplasmic [26]. Using over-expression and RNAi experiments, Chiorazzi *et al.* demonstrated that FBXO10 is the BCL2-binding subunit of an SCF cytoplasmic ubiquitin ligase complex that ubiquitinates BCL2 to trigger proteasomal degradation in DLBCL. The relevance of this process was supported by infrequent *FBXO10* partial loss-of-function somatic mutations and frequently reduced mRNA expression in DLBCL samples from their cohort [26]. Low *FBXO10* mRNA resulting in high BCL2 also appears to drive accumulation of mantle cell lymphomas (MCL) [29] derived from marginal zone or memory B cells [30].

Mice expressing *BCL2* under the control of the *IGH* enhancer (Eμ) have increased accumulation of BCL2 protein in B cells, dramatically increased numbers of mature B cells and GC B cells, and develop low-incidence pre-B lymphomas, immunoblastic lymphomas and plasmacytomas [31–34]. Constitutive over-expression of BCL2 in all hematopoietic lineages, in transgenic mice where the human *BCL2* gene is fused to the *Vav* gene promoter, has a potent effect on the survival, development and maturation of many blood cell types [35] and results in increased incidence of follicular lymphoma [36]. We therefore hypothesized that mice with germline *Fbxo10* loss-of-function mutations would have increased BCL2 protein in B cells and correspondingly increased B cell and GC B cell accumulation, and increased BCL2 and dysregulated survival in other blood cell types. Here, we tested this hypothesis by analysing mice with either a CRISPR/*Cas9*-engineered germline deletion in *Fbxo10* or partial loss-of-function E54K missense mutation in the F-box domain.

## Results

### Germline *Fbxo10*^*E54K*^ and *Fbxo10*^*frameshift*^ mutant mice appear at Mendelian frequencies, present with no visible clinical phenotype and age normally

Our interest in *FBXO10* was stimulated by the identification of a very rare, predicted damaging, missense variant E54K inherited in homozygous state from healthy heterozygous parents in a child with multiple autoimmune diseases and possible learning difficulties (unpublished data). We have since discovered compound heterozygous *TNFAIP3* mutations explaining the child’s autoimmunity, but it was notable that E54K had been independently found as a heterozygous *de novo* mutation in a child with autism spectrum disorder [37]. The E54 residue lies within the F-box superfamily domain (SSF81383, FBXO10 residues 6-80) required for SCF complex assembly, is strictly conserved from fish to humans, and the substitution from glutamic acid to lysine represents a non-conservative charge reversal (Fig 1A,B). When FLAG-tagged FBXO10 was expressed in HEK293T cells, the E54K substitution decreased immunoprecipitation of endogenous SKP1 to a similar extent as the partial loss-of-function R44H FBOX mutation (Fig 1C) previously characterised in a human lymphoma [26].

**Fig 1.**
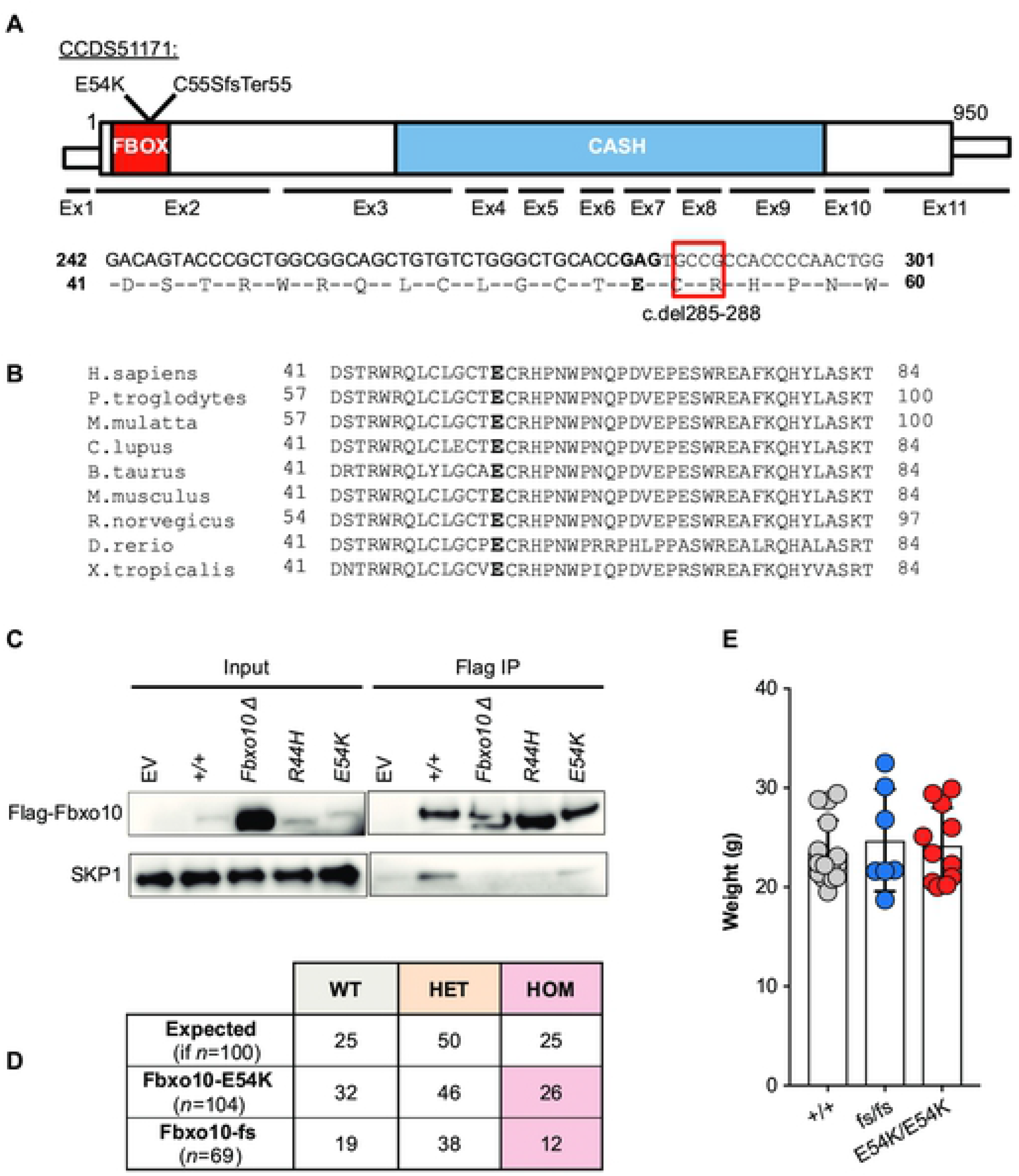
Viable mice with homozygous germline *Fbxo*^*10E54K*^ or *Fbxo10*^*fs*^ mutations. **(A)** Schematic of mouse *Fbxo10* mRNA CCDS51171, showing position of exons, location of mutations, FBOX (SSF81383) and three tandem CASH (SM00722) domains, and the four cDNA nucleotides deleted in the *Fbxo10*^*C55SfsTer55*^ (*fs*) allele. **(B)** Alignment of the FBOX10 amino acid sequence from the indicated species: E54 in bold. **(C)** Expression vectors, either empty or encoding FLAG-tagged human FBXO10 wildtype or with the indicated mutations, were transfected into HEK293T cells and lysates or anti-FLAG immunoprecipitates western blotted with antibodies to FLAG or SKP1. **(D)** Expected and observed numbers of offspring of the indicated genotypes from intercrossed heterozygous parents. Statistical analysis by Chi-Square test with *n* = 2 degrees of freedom, testing for differences relative to a 1WT:2HET:1HOM expected Mendelian ratio (p = 0.61 and p = 0.55 for *Fbxo10*^*E54K*^ or *Fbxo10*^*fs*^ respectively). **(E)** Body weight of *Fbxo10*^+/+^, *Fbxo10*^*fs/fs*^ and *Fbxo10*^*E54K/E54K*^ mice 9-20 weeks old (p = 0.51 and p = 0.62 for *Fbxo10*^*E54K*^ or *Fbxo10*^*fs*^, respectively). Each dot represents an individual mouse of the indicated genotype. Statistical comparison between each mutant and wild-type group was performed by t-test corrected for multiple comparisons using the Holm-Sidak method.

To explore E54K as a candidate mutation, *Fbxo10*^*E54K*^ mice were produced by CRISPR/*Cas9* gene editing in mouse embryos following established molecular and animal husbandry techniques [38]. Two independent alleles were engineered and propagated in C57BL/6J mice (Fig 1A): a point mutation in exon 2 changing the Glutamate 54 codon to Lysine (E54K), or a 4 nucleotide deletion in codons 55 and 56 within exon 2 changing the Cysteine 55 codon to Serine and creating a reading frame shift and premature stop codon after 55 codons (c.del285_288 or p.Cys55SerfsTer55; abbreviated as *fs*). The *fs* deletion does not create a new splice donor site and there is no evidence of alternate splice forms of *Fbxo10* that skip exon 2 in mouse or human. It is therefore likely to create a null allele, although we lack suitable antibodies to test for a protein remnant in primary mouse cells. When heterozygous animals were intercrossed, neither E54K nor the frameshift mutation resulted in altered frequencies of heterozygous or homozygous mutant mice relative to expected Mendelian ratios (Fig 1D). Adult homozygous mutant animals up to 50 weeks old appeared normal and healthy, and had no significant difference in body weight from wild-type littermates (Fig 1E).

### FBXO10 deletion or missense mutation had no detectable effect on B cells in bone marrow or spleen

Flow cytometric analysis of early B cell development in the bone marrow of adult 10-20 week old mice (data not shown) and 40-50 week old mice (Fig 2A,B) revealed no discernable difference between *Fbxo10* wild-type and mutant mice in the frequencies of bone marrow B cells, nor in the subsets of mature recirculating B cells, immature B cells, or the different stages of precursor B cell differentiation. Congruent with this, there were no detectable differences in expression of B220, CD19, CD93, IgM, IgD, CD21, CD23 or CD86 in these various subsets (S1 Fig A).

**Fig 2.**
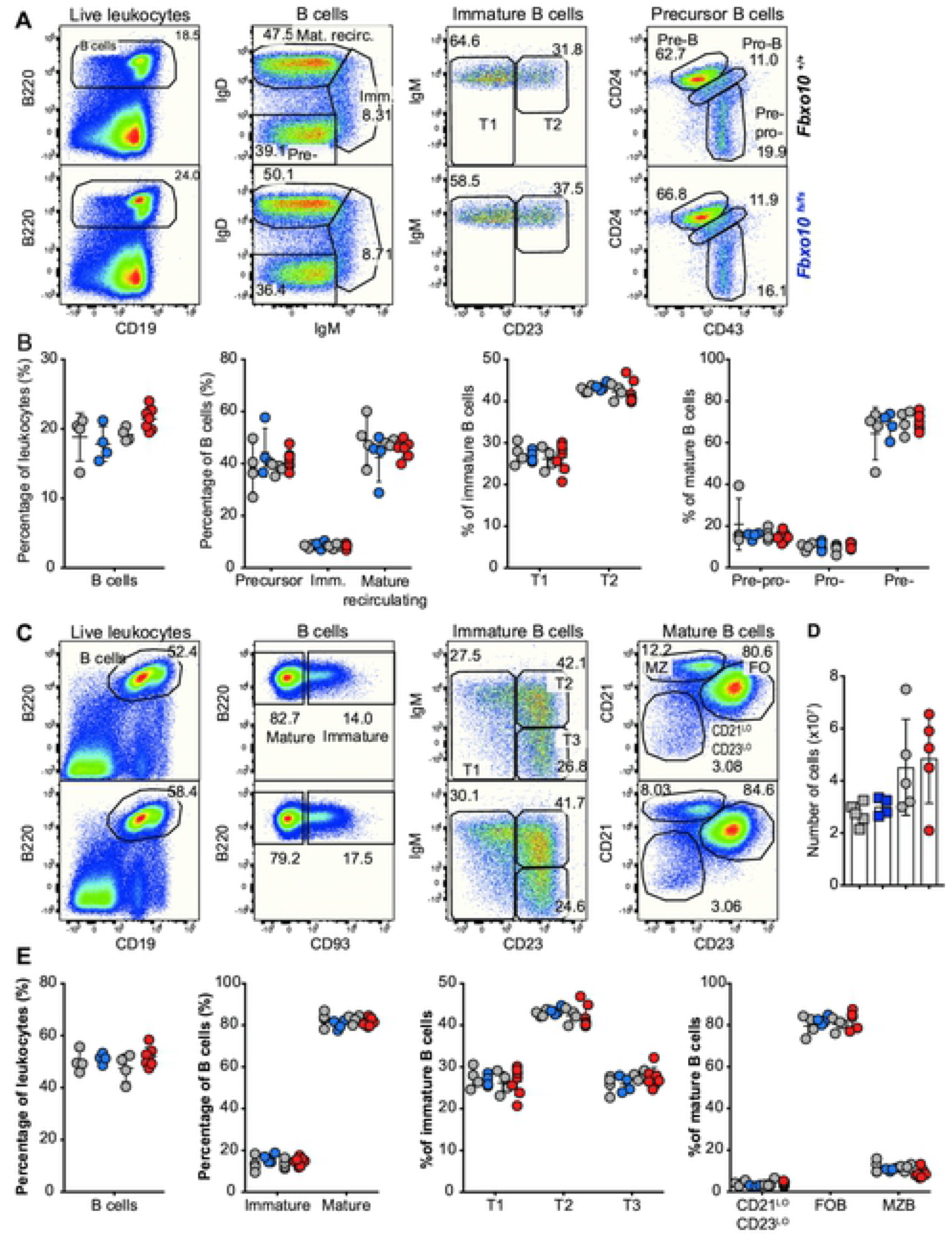
*Fbxo10* frameshift or missense mutation do not discernably affect B cell subsets in the bone marrow or spleen. **(A)** Representative flow cytometric gating strategy to delineate B cell developmental subsets in the bone marrow. Numbers denote cells in the gate as percentage of parent population. **(B)** Frequency of indicated B cell subsets in the bone marrow in *Fbxo10*^*fs/fs*^ and *Fbxo10*^+/+^ littermate control mice (blue and left set of grey circles, respectively) and in *Fbxo10E54K/$54K* and *Fbxo10+/+* littermate control mice (red and right-hand grey circles, respectively). **(C)** Representative flow cytometric gating strategy to delineate splenic B cell subsets. **(D)** Spleen cellularity in *Fbxo10*^*fs/fs*^ and littermate control mice (blue and grey circles, respectively) and in *Fbxo*^*10E54K/$54K*^ and *Fbxo10+/+* littermate control mice (red and grey circles, respectively). **(E)** B cell subsets in the spleen of *Fbxo10*^+/+^, *Fbxo10*^*fs/fs*^ and *Fbxo10*^*E54K/E54K*^ mice. **(B, D, E)** Each dot represents data from an individual animal. Data are representative of *n* = 2 experiments on mice 40-50 weeks old, and similar results observed in *n* = 2 experiments on mice 10-20 weeks old. Statistical analysis: t-test corrected for multiple comparisons using the Holm-Sidak method yielded no evidence for significant differences between mutants and wild-type controls with p < 0.05.

Spleen cellularity was also unaffected (Fig 2D) and there was no sign of lymphadenopathy in the mutant mice (data not shown). Neither mutation resulted in any changes in distribution of B cell maturation subsets in the spleen of 40-50 week old (Fig 2C,D) or 10-12 week old mice (data not shown), nor in any detectable changes in surface expression by splenic B cell subsets of the markers listed above (S1 Fig B). Once more, the lack of a visible effect of *Fbxo10* mutation or deletion, even in elderly mice, indicates that FBXO10 plays no role or a functionally redundant role in B cell early development and splenic B cell maturation.

### FBXO10 deletion or missense mutation had no visible effect on the magnitude or quality of a polyclonal GC B cell response to SRBC immunisation

FBXO10 is particularly highly expressed by GC B cells and appears to be most important for regulating BCL2 protein levels in GCB-type DLBCL, based on the reduced *FBXO10* mRNA expression and low frequency heterozygous *FBXO10* hypomorphic missense alleles in DLBCL and the high *FBXO10* mRNA expression in GC B cells [26]. We therefore tested for increased accumulation of GC B cells in *Fbxo*^*10E54K*^ and *Fbxo10*^*fs*^ mice in a T cell-dependent response following sheep red blood cell (SRBC) immunisation. Sacrifice of *Fbxo10*^*fs/fs*^, *Fbxo10*^*E54K/E54K*^ and wild-type mice 7 days post SRBC-immunisation demonstrated no significant difference in the magnitude of the GC response, nor the dark zone/light zone distribution or the fraction of IgG1 class-switched GC B cells (Fig 3A,B).

**Fig 3.**
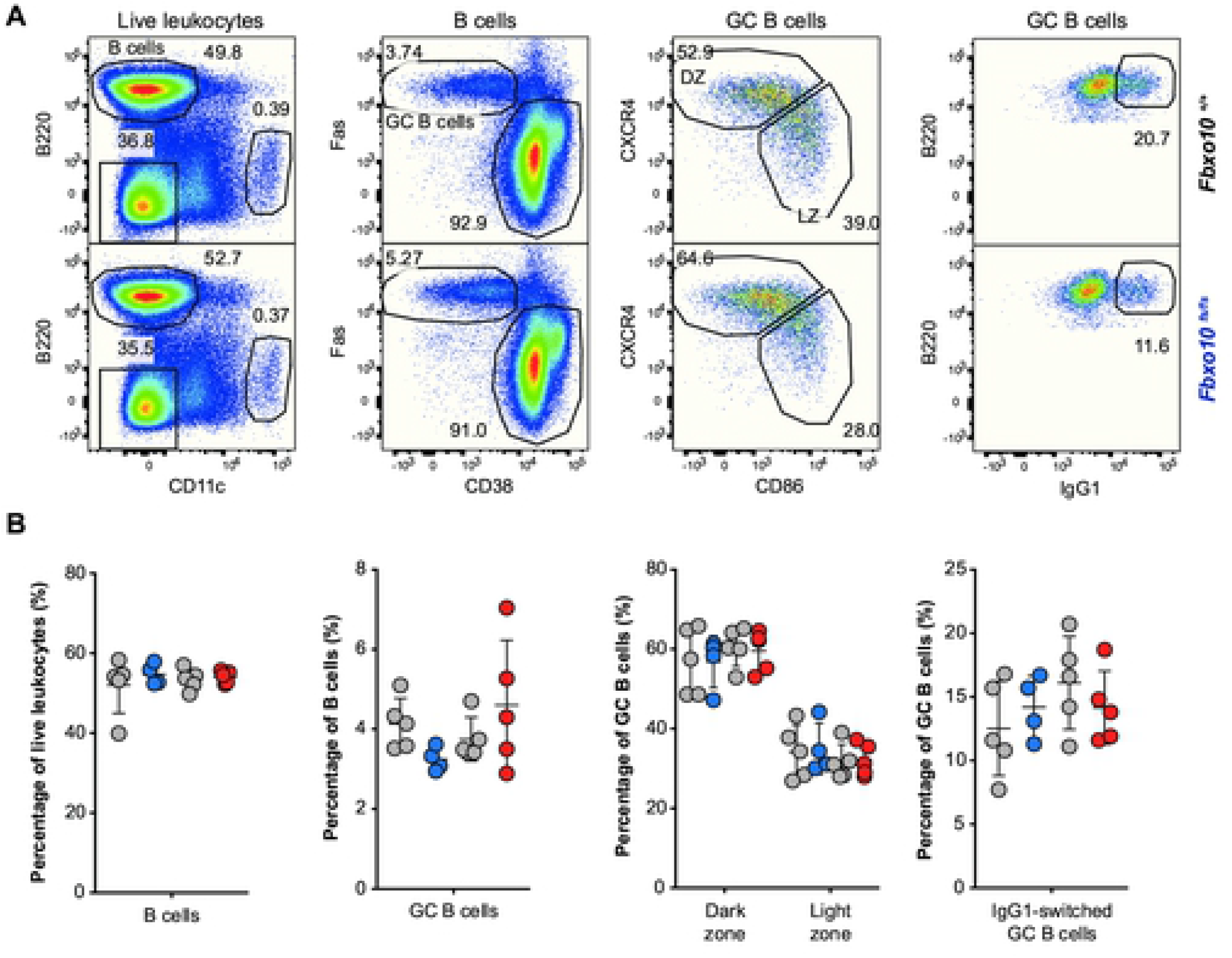
*Fbxo10* frameshift or missense mutation do not visibly expand or alter the GC response to SRBC immunisation. **(A)** Representative flow cytometric gating strategy to delineate GC B cells responding to SRBC immunisation. Numbers denote cells in the gate as a percentage of parent population. **(B)** Frequency of B cells, GC B cells, light zone/dark zone distribution and IgG1-class-switched GC B cells in the spleen of *Fbxo10*^+/+^ (grey circles matched with mutant siblings) *Fbxo10*^*fs/fs*^ (blue circles), and *Fbxo10*^*E54K/E54K*^ (red circles) mice day 7 post-immunisation. B: each dot represents an individual biological replicate. Data are representative of *n* = 2 experiments on mice 10-20 weeks old. Statistical analysis: t-test corrected for multiple comparisons using the Holm-Sidak method yielded no evidence for significant differences between mutants and wildtype controls with p < 0.05.

### FBXO10 deletion or missense mutation had no visible effect on B or T cell expression of putative FBXO10 targets

Given the lack of any increase in mature B cells or GC B cells, as would be expected if FBXO10 deficiency resulted in increased BCL2 protein accumulation, we measured BCL2 protein levels in single B cells by intracellular antibody staining followed by flow cytometric analysis. Interestingly, we observed no differences in expression of BCL2 between wild-type and mutant GC B cells (Fig 4A,B) The same was true for BCL6 (Fig 4C,D) and BAFF-R that is an FBXO11 target in lymphoma (Fig 4E,F). The same was true for non-GC B cells as well as for effector memory, central memory and naïve CD4 and CD8 T cells (Fig 4A-F). Importantly, the well-validated changes in expression of BAFF-R, BCL-6, and BCL-2 between lymphoid subsets, such as increased BCL-6 expression in GC B cells relative to non-GC B cells or increased BCL-2 expression in effector memory T cells relative to naïve T cells, provided a useful internal control to validate successful staining in terms both of specificity and sensitivity (Fig 4B,D,F).

**Fig 4.**
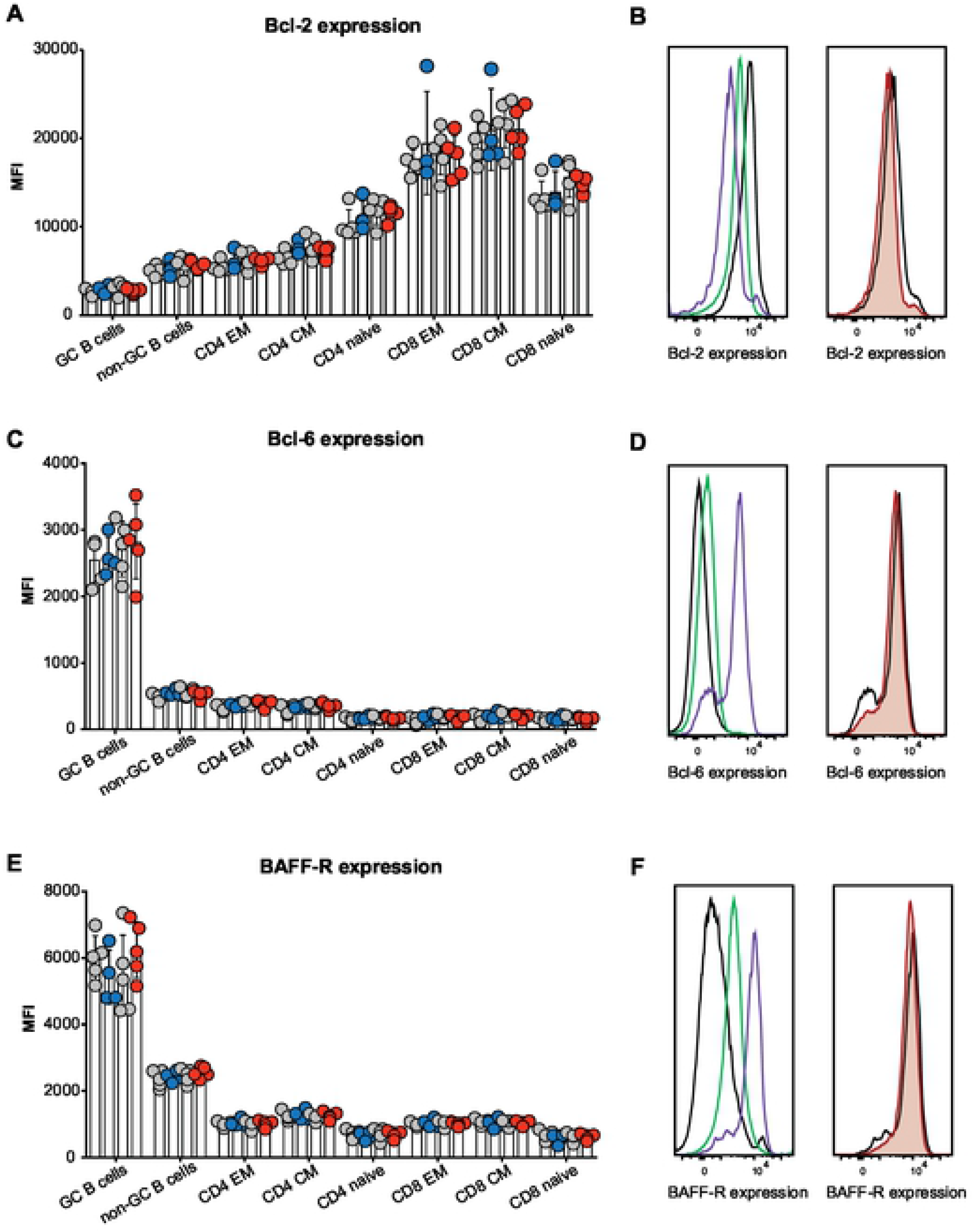
*Fbxo10* frameshift or missense mutation do not discernably alter the protein expression levels of putative Fbxo10 targets in lymphoma. **(A,C,E)** Mean fluorescence intensity (MFI) for BCL2, BCL6 or BAFF-R expression, respectively, in splenic B and T cell subsets of *Fbxo10*^+/+^, *Fbxo10*^*fs/fs*^, *Fbxo10*^*E54K/E54K*^ mice 10-20 weeks old, day 7 post-immunisation with SRBCs. **(B,D,F)** Left panel: representative histogram overlay of fluorescent antibody staining for intracellular BCL-2 or BCL-6 or cell surface BAFF-R in T cells (black), non-GC B cells (green) or GC B cells (purple). Right panel: representative histogram overlay for BCL2, BCL6 or BAFF-R expression in *Fbxo10*^+/+^ (black) versus *Fbxo10*^*E54K/E54K*^ (red) GC B cells. **E**ach dot represents an individual animal, with genotypes as in Figure 2. Data are representative of *n* = 2 experiments on mice 10-20 weeks old, 7 days post-immunisation with SRBC, and similar results obtained for *n* = 2 experiments on un-immunised mice 40-50 weeks old. Statistical analysis: t-test corrected for multiple comparisons using the Holm-Sidak method yielded no evidence for significant differences between mutants and wildtype controls with p < 0.05.

### FBXO10 deletion or missense mutation had no detectable effect on T cell thymic development or T cell splenic maturation

Because expression of the *Vav-BCL2* transgene in mice also causes a marked elevation of T lymphocytes and altered relative abundances of developing CD4− CD8− double negative (DN), CD4+ CD8+ double positive (DP) and single positive (SP) thymocytes [35], we investigated T cell development and maturation in *Fbxo*^*10E54K*^ or *Fbxo10*^*fs*^ mutant mice. Our analysis revealed no significant difference in fractions of thymic DN, DP, CD4 and CD8 SP T cells in elderly (40-50 week old) *Fbxo10*^*fs*^ and *Fbxo*^*10E54K*^ mutant mice, relative to their wild-type littermate controls, nor in the fractions of early developing DN1-DN4 thymocytes (Fig 5A,B). There were also no detectable changes in expression of CD25, CD44, CD69, PD1 and CD62L by these thymic subsets (S2 Fig A,B). In our hands, the only significant effect of *Fbxo10* deletion or missense mutation was a very slight increase in frequency of Tregs in the thymus (Fig 5A,B) that was a consistent trend in different cohorts. Splenic T cell subsets were also not significantly affected by *Fbxo10* mutations, as the percentage of T cells, CD4:CD8 ratio, fraction of Tregs and of CD4 and CD8 effector memory, central memory and naïve subsets were comparable in mutant relative to wild-type mice (Fig 5C,D). Similarly, no changes in expression of maturation/activation markers were detected in these various subsets between wild-type and mutant mice (S2 Fig C).

**Fig 5.**
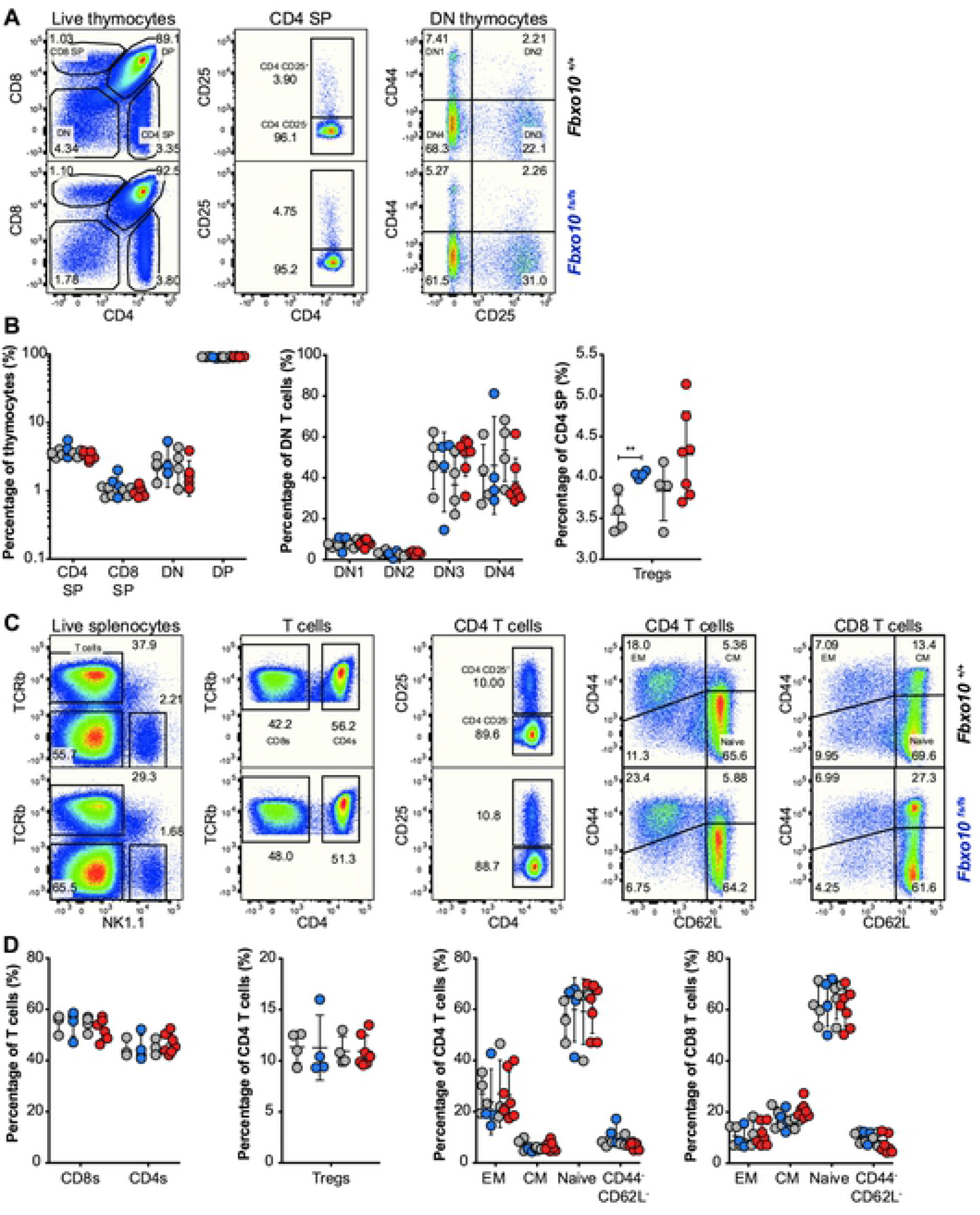
*Fbxo10* frameshift or missense mutation do not discernably alter thymic or spleen T cell subsets. **(A)** Representative flow cytometric gating strategy to delineate T cell developmental populations in the thymus. Numbers denote cells in gate as percentage of parent population. **(B)** T cell developmental subsets in the thymus of *Fbxo10*^+/+^, *Fbxo10*^*fs/fs*^, *Fbxo10*^*E54K/E54K*^ mice. **(C)** Representative flow cytometric gating strategy to delineate splenic T cell subsets. **(D)** T cell subsets in the spleen of *Fbxo10*^+/+^, *Fbxo10*^*fs/fs*^, *Fbxo10*^*E54K/E54K*^ mice. B, D: each dot represents data from an individual mouse: *Fbxo10*^+/+^ in grey, *Fbxo10*^*fs/fs*^ in blue, *Fbxo10*^*E54K/E54K*^ in red as in Figure 2. Results representative of *n* = 2 experiments on mice 40-50 weeks old, and similar results observed in *n* = 2 experiments on mice 10-20 weeks old. Statistical analysis: t-test corrected for multiple comparisons using the Holm-Sidak method, ** p < 0.01, all other differences were not significant.

This was also true for young *Fbxo10* mutant mice (data not shown), and the lack of any observable effects even in elderly mice, despite the common exacerbation of underlying immune defects with age in mice and humans, indicates that FBXO10 plays no or a redundant role in murine T cell development and maturation, at least in un-immunised mice. Further analysis using antigen-specific challenge models may reveal a context-specific role for FBXO10 in T cells. *Fbxo10* expression has for example been shown to increase in Jurkat cells upon cellular stress, downstream of LEDGF signalling [39]. Nevertheless, we can conclude that FBXO10 alone is not required for the development, differentiation or survival of T cells in mice.

### FBXO10 deletion or missense mutation had no detectable effect on the distribution of murine lymphoid and myeloid leukocyte subsets in the bone marrow or spleen

Similarly, despite the changes in lymphoid and myeloid subsets in *Vav-BCL2* transgenic mice [35], the distribution of NK cells, dendritic cells, monocytes/macrophages and dendritic cells in the bone marrow (Fig 6A,C) and spleen (Fig 6B,D) was not significantly different between *Fbxo10* mutant or wild-type mice 10-12 weeks old (data not shown) or 40-50 weeks old (Fig 6A-D). No consistent changes in size, granularity or expression of CD11b, CD11c, Ly6G, Ly6C, CD44, CD62L proteins were detected in any leukocyte subset in the bone marrow (S3 Fig A) or spleen (S3 Fig B) of *Fbxo10*-mutant mice. As many of these proteins are well-validated markers of activation and differentiation of myeloid cells, we may infer that development and maturation of myeloid cells are largely unaffected by *Fbxo*^*10E54K*^ and *Fbxo10*^*fs*^.

**Fig 6.**
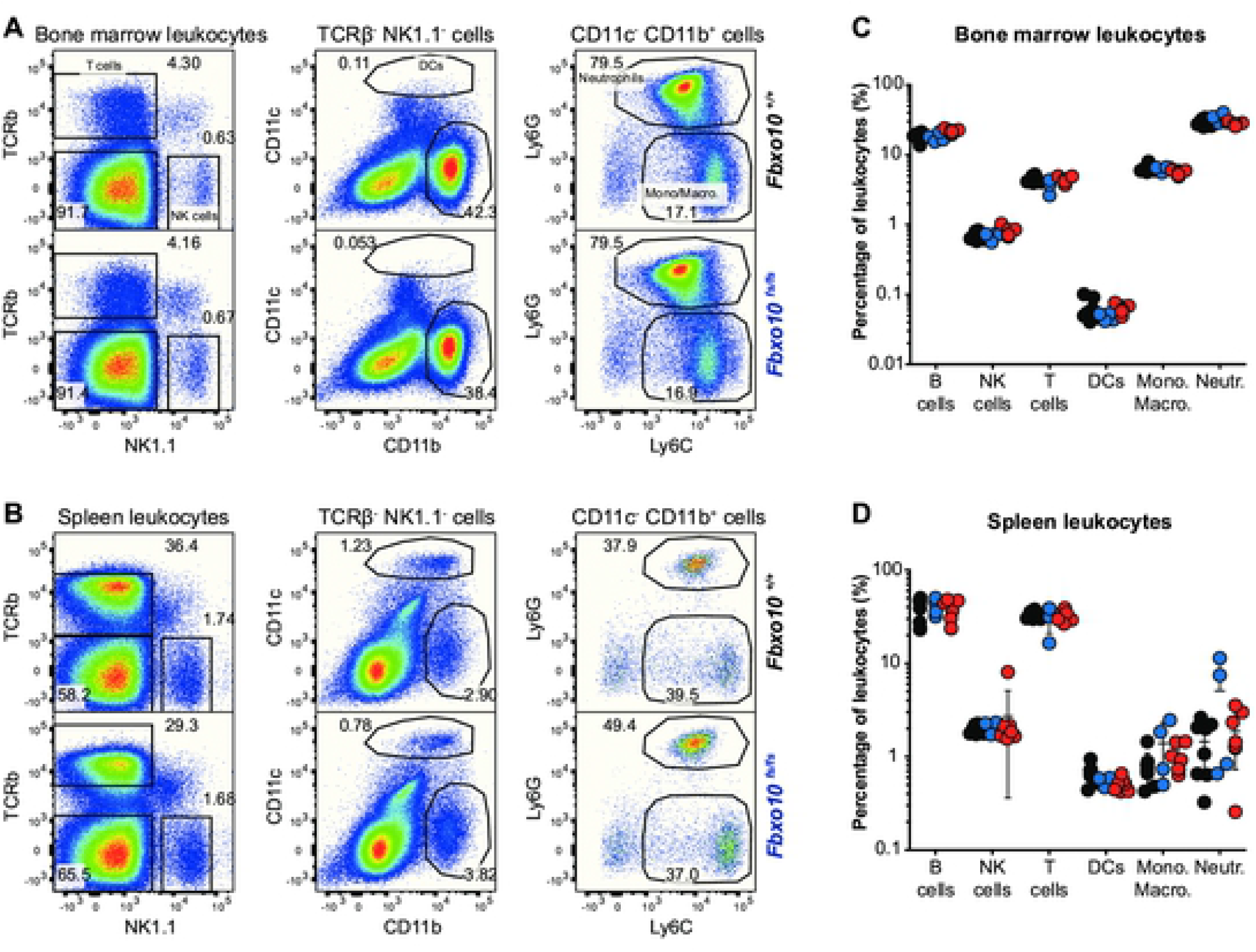
*Fbxo10* frameshift or missense mutation do not cause discernable differences in myeloid and lymphoid leukocytes within the spleen or bone marrow. **(A)** Representative flow cytometry gating strategy to delineate leukocyte populations in the bone marrow. Numbers denote cells in gate as percentage of parent population. **(B)** Representative flow cytometric gating strategy to delineate leukocyte populations in the spleen. **(C)** leukocyte subsets in the bone marrow of *Fbxo10*^+/+^, *Fbxo10*^*fs/fs*^, *Fbxo10*^*E54K/E54K*^ mice. **(D)** leukocyte subsets in the spleen of *Fbxo10*^+/+^, *Fbxo10*^*fs/fs*^, *Fbxo10*^*E54K/E54K*^ mice. C, D: each dot represents an individual biological replicate. Results representative of *n* = 2 experiments on mice 40-50 weeks old, and similar results observed in *n* = 2 experiments on mice 10-20 weeks old. *Fbxo10*^+/+^ in black, *Fbxo10*^*fs/fs*^ in blue, *Fbxo10*^*E54K/E54K*^ in red. Statistical analysis: t-test corrected for multiple comparisons using the Holm-Sidak method yielded no evidence for significant differences between mutants and wildtype controls with p < 0.05.

## Discussion

Based on somatic mutations in DLBCL, germline mutations in *C elegans*, and experimental overexpression and RNAi knockdown experiments in human DLBCL cells, we hypothesised that germline loss of function *Fbxo10* mutations in mice would cause increased BCL2 protein accumulation in mature B cells and GC B cells and corresponding increased B cell accumulation. This hypothesis was not supported here by characterisation of C57BL/6J mice with a germline missense mutation or frameshift mutation in *Fbxo10.* No visible morphological or immune cellular phenotype resulted from FBXO10 loss-of-function in mice. Mutant mice presented with normal breeding frequencies (Fig 1), spleen cellularity (Fig 2), unchanged B and T cell early development and splenic maturation (Fig 2 and 5), normal frequencies of leukocyte subsets in the bone marrow or spleen (Fig 6), and identical expression of protein markers associated with development, differentiation, activation and migration in all of these various leukocyte subsets (S1-S3 Fig).

Mutation or loss of *Fbxo10* did not affect the magnitude of a polyclonal GC B cell response to SRBC immunisation, nor the GC dark zone/light zone distribution or frequency of IgG1 class-switched cells (Fig 3). Finally, wild-type and mutant lymphoid subsets presented with identical expression of BCL2 (Fig 4). These observations were made not only in 10-12 week old adult mice, but also in elderly 40-50 week old mice where one can often observe exacerbation of underlying immune defects over time. Of 59 mice aged to 30-50 weeks old, no *Fbxo10*-mutant (or wild-type) mouse developed a solid organ or lymphoid malignancy. Thus, *Fbxo10* hypomorphic mutation or deletion results in no visible changes in expression of FBXO10 target BCL2 in mice, even in the GC (nor of FBXO11 target BCL6). The importance of FBXO10 function to BCL2 expression and survival of DLBCL [26, 28] may be associated with a concomitant loss of redundant or compensatory mechanisms in these cells.

Another role identified for FBXO10 in human lymphoma cell lines is in the negative regulation of BCR signalling via BCR signalling-induced membrane re-localisation followed by degradation of human germinal-centre associated lymphoma (HGAL, also called GCET2) protein levels [41]. HGAL is GC B cell-specific, enhances BCR signalling by increasing activation of Syk downstream effectors and human *HGAL*-transgenic mice develop lymphoid hyperplasia in elderly mice [42]. Notably however, deletion of *HGAL* (also called *M17*) had no effect on the GC response in mice [43]. The normal GC responses observed in mice with frameshift or missense FBXO10 do not support a critical role for FBXO10 in degrading HGAL in mice, although we have not measured HGAL levels in mutant GC B cells.

The related protein, FBXO11, may theoretically compensate for FBXO10 loss of function mutations. In the gnoMAD database analysing 124,000 adult human exomes or genomes, FBXO11 has a pLI=1.0, due to much lower than expected occurrence of heterozygous stop gain or frameshift mutations. This is consistent with evidence for human FBXO11 haploinsufficiency, with heterozygous germline *de novo* loss-of-function alleles found recurrently in children with neurodevelopmental disorders [37, 44, 45], and with high frequency heterozygous loss-of-function somatic mutations in human B cell lymphoma [28]. In mice, an *Fbxo11* missense mutation in the CASH domain causes a heterozygous developmental disorder of the ear and homozygous lethal dysmorphism [46], while homozygous conditional *Fbxo11* deletion in GC B cells increases their number and BCL6 protein levels [47]. By contrast, FBXO10 has a pLI=0 in gnoMAD indicating that heterozygous null mutations occur at the expected frequency in the adult human population. Evidence against FBXO11 as a redundant paralogue for BCL2 regulation comes from FBXO11 siRNA knockdown in human B cell lymphoma cells, which dramatically enhanced BCL6 protein stability after protein translation was pharmacologically blocked, but did not enhance BCL2 protein stability analysed in the same cell lysates [28].

Another candidate compensatory BCL2-regulator is ARTS (gene name *SEPT4*), which serves as an adapter to promote BCL2 ubiquitination by the XIAP ubiquitin ligase in apoptotic cells [48]. ARTS-deficient B cells in mice nevertheless develop and accumulate in normal numbers, suggesting that ARTS is also unnecessary or redundant for regulating BCL2-dependent B cell survival [49]. However, ARTS deficiency does promote exaggerated mature B cell accumulation in Emu-MYC transgenic B cells where MYC is dysregulated and promotes apoptosis, and this effect is abolished in ARTS-XIAP double-deficient B cells [49]. Given the importance of balanced BCL2 and BIM protein levels for controlling normal B cell survival and suppressing B cell lymphoma [1], it would not be surprising that BCL2 protein turnover be governed by multiple, redundant ubiquitin ligases.

To our knowledge, our results constitute the first characterisation of mice with homozygous loss-of-function mutations in FBXO10. They highlight the importance of investigating the functional redundancy/synergy of FBXO10 loss-of-function with mutations in other pathways and with loss-of-function of SCF complex members such as FBXO11. The incongruity between the role of FBXO10 in inducing cell death of the *C. elegans* tail spike cell and of human DLBCL cells relative to leukocyte subsets in the mouse should be further investigated.

## Materials and Methods

### Mice

Mice were bred at Australian BioResources (MossVale, NSW, Australia) and kept in specific pathogen-free conditions at the Garvan Institute (Sydney, Australia). All animal studies were approved and conducted in compliance with the guidelines set by the Garvan/St.Vincent’s Animal Ethics Committee.

*Fbxo10*^*E54K*^ and *Fbxo10*^*KO*^ mice were produced by the Mouse Engineering Garvan/ABR (MEGA) Facility using CRISPR/Cas9 gene targeting in mouse embryos following established molecular and animal husbandry techniques (Yang et al., 2014). The single guide RNA (sgRNA) was based on a target site exon 2 of Fbxo10 (CCAGTTGGGGTGGCGGCACTCGG) (protospacer-associated motif = PAM italicised and underlined) and was microinjected into the nucleus and cytoplasm of C57BL/6J zygotes together with polyadenylated S.pyogenes Cas9 mRNA and a 150 base single-stranded, anti-sense, deoxy-oligonucleotide homologous recombination substrate carrying the E54K (GAG>AAG) mutation and a PAM-inactivating silent mutation in the T53 codon (ACC>ACA). A founder mouse heterozygous for both substitutions was obtained and used to establish the Fbxo10E54K line. An additional founder carrying a 4bp frame shift mutation after the first base of the C55 codon was bred to establish the Fbxo10⍼ line. Both lines were maintained on an inbred C57BL/6J background. All experiments were approved by the Garvan/St Vincent’s Animal Ethics Committee. Mice were bred and housed in specific pathogen-free conditions at Australian BioResources (Moss Vale) and the Garvan Institute Biological Testing Facility.

### Flow cytometric analysis

Mouse organs were harvested into FACS buffer (PBS/1% BSA/0.02% sodium azide) and single cell suspensions passed through a 70 μm cell strainer (Falcon, Corning, NY, USA). In analysis of spleen or blood immune subsets, red blood cell (RBC) lysis was performed using lysis buffer solution (0.8% ammonium chloride, 0.08% sodium bicarbonate, 0.04% EDTA disodium salt, pH 7.3).

Single cell suspensions were stained with antibodies targeting cell-surface (B220, BAFF-R, IgM, IgD, IgG1, Ly-6C, Ly-6G, PD-1, TCRβ, CD3, CD11b, CD11c, CD19, CD21/35, CD23, CD24, CD25, CD28, CD38, CD43, CD44, CD62L, CD69, CD86, CD93, CD95, CD278) or intracellular proteins (BCL-2, BCL-6, CD152) and cells were acquired on an LSR II analyser (BD Pharmingen), followed by flow cytometric analysis using the FlowJo Software (FlowJo LLC, Ashland, OR, USA).

### Immunoprecipitation

Expression vectors, either empty or encoding FLAG-tagged human FBXO10 wild-type or with the indicated mutations, were transfected into HEK293T cells and lysates or anti-FLAG immunoprecipitates western blotted with antibodies to FLAG or SKP1. Immunoprecipitations were performed as previously described [26], using FLAG (Sigma F3165) and SKP1 (Santa Cruz sc-5281) antibodies. Construction of cDNA of FLAG-tagged FBXO10, ΔFBXO10 and FBXO10 R44H in a retroviral vector and transfection into HEK293T cells were also performed as previously described [26]. FBXO10 E54K was generated by site-directed mutagenesis (Stratagene 200521-5) using the following primers:

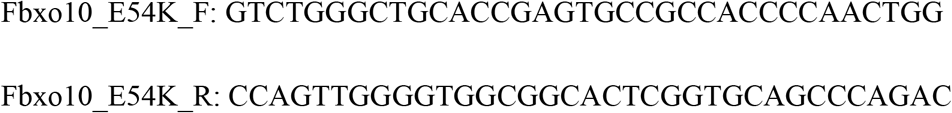

### Statistical analysis

GraphPad Prism 6 (GraphPad Software, San Diego, CA, USA) was used for analysis of flow cytometry or ELISA data. For comparisons between genotypes, the variance was approximately equal between samples and comparisons were made using a Student’s t-test, and corrected for multiple comparisons using the Holm-Sidak method. For these tests, p < 0.05 was considered statistically significant. In all flow cytometry summary figures, each data point represents an individual mouse. Error bars indicate the mean and standard distribution. *p<0.05; **p<0.01; ***p<0.001.

## Acknowledgements

We thank the Garvan Institute ABR, GMG and Flow Cytometry facilities for expert animal husbandry, genotyping and cell sorting. This work was supported by NHMRC Grants APP1113904, APP1081858, and APP1108800 and by the Ritchie Family Foundation.

## Supporting information

**S1 Fig. Representative graphs showing that Fbxo10 deletion or mutation do not alter the protein expression of B cell development and maturation maturation markers in bone marrow or spleen.**

**(A)** Mean fluorescence intensity (MFI) for IgM, IgD, CD93, CD86 expression in splenic B cell subsets of *Fbxo10*^+/+^, *Fbxo10*^*fs/fs*^, *Fbxo10*^*E54K/E54K*^ mice 40-50 weeks old. **(B)** MFI for IgM, IgD, CD24, CD43 expression in bone marrow B cell subsets of *Fbxo10*^+/+^, *Fbxo10*^*fs/fs*^, *Fbxo10*^*E54K/E54K*^ mice 40-50 weeks old. Each dot represents an individual biological replicate in *Fbxo10*^+/+^ (black), *Fbxo10*^*E54K/E54K*^ (red) or *Fbxo10*^*fs/fs*^ (blue) mice. Similar results were obtained for multiple protein markers (CD19, CD21/35, CD23, CD24, CD43, CD86, etc.). Results are representative of *n* = 2 experiments on un-immunised mice 40-50 weeks old and similar results were obtained for *n* = 2 experiments on mice 10-20 weeks old, 7 days post-immunisation with SRBC. Statistical analysis: t-test corrected for multiple comparisons using the Holm-Sidak method yielded no evidence for significant differences between mutants and wildtype controls with p < 0.05.

**S2 Fig. Representative plots showing that Fbxo10 deletion does not alter the protein expression levels of T cell activation and maturation markers in thymus or spleen. (A,B)**

Representative histogram overlays of *Fbxo10*^+/+^ (grey fill) or *Fbxo10*^*fs/fs*^ (blue fill) thymocyte subsets relative to *Fbxo10*^+/+^ control thymocytes (black line) for CD44 (A) or for CD69 (B). Results are representative of results obtained for other markers: CD25, CD69, PD1, CD3, etc. **(C)** Representative histogram overlays of *Fbxo10*^+/+^ (grey fill) or *Fbxo10*^*fs/fs*^ (blue fill) splenic T cells relative to *Fbxo10*^+/+^ control cells (black line) showing CD62L (left 3 panels) or CD44 (right 3 panels). Results are representative of results obtained for other markers: CD25, CD62L, PD1, CD3, etc.

**S3 Fig. Representative graphs showing that Fbxo10 deletion or mutation do not alter the protein expression of development and maturation markers for leukocytes in bone marrow or spleen.**

**(A)** Mean fluorescence intensity (MFI) for CD11b, Ly6G, CD62L, CD44 expression in spleen leukocyte subsets of *Fbxo10*^+/+^, *Fbxo10*^*fs/fs*^, *Fbxo10*^*E54K/E54K*^ mice 40-50 weeks old. **(B)** Mean fluorescence intensity (MFI) for CD11b, Ly6G, CD62L, CD44 expression in bone marrow leukocyte subsets of *Fbxo10*^+/+^, *Fbxo10*^*fs/fs*^, *Fbxo10*^*E54K/E54K*^ mice 40-50 weeks old. Each dot represents an individual biological replicate in *Fbxo10*^+/+^ (black), *Fbxo10*^*E54K/E54K*^ (red) or *Fbxo10*^*fs/fs*^ (blue) mice. Similar results were obtained for multiple protein markers (NK1.1, Ly6G, FSC, SSC-A, MHC II, etc.). Results are representative of *n* = 2 experiments on un-immunised mice 40-50 weeks old and similar results were obtained for *n* = 2 experiments on mice 10-20 weeks old, 7 days post-immunisation with SRBC. Statistical analysis: t-test corrected for multiple comparisons using the Holm-Sidak method yielded no evidence for significant differences between mutants and wildtype controls with p < 0.05.

